# Mononuclear cell dynamics in *M. tuberculosis* infection provide opportunities for therapeutic intervention

**DOI:** 10.1101/347294

**Authors:** Brian A. Norris, Joel D. Ernst

## Abstract

*Mycobacterium tuberculosis* causes chronic infection of mononuclear phagocytes, especially resident (alveolar) macrophages, recruited macrophages, and dendritic cells. Despite the importance of these cells in tuberculosis (TB) pathogenesis and immunity, little is known about the population dynamics of these cells at the sites of infection. We used a combination of congenic monocyte adoptive transfer, and pulse-chase labeling of DNA, to determine the kinetics and characteristics of trafficking, differentiation, and infection of mononuclear phagocytes during the chronic, adaptive immune phase of *M. tuberculosis* infection in mice. We found that Ly6C^hi^ monocytes traffic rapidly to the lungs, where a subpopulation become Ly6C^lo^ and remain in the lung vascular space, while the remainder migrate into the lung parenchyma and differentiate into Ly6C^hi^ dendritic cells, CD11b^+^ dendritic cells, and recruited macrophages. As in humans with TB, *M. tuberculosis*-infected mice have increased numbers of blood monocytes; this is due to increased egress from the bone marrow, and not delayed egress from the blood. Pulse-chase labeling of dividing cells and flow cytometry analysis revealed a T_1/2_ of ∼15 hrs for Ly6C^hi^ monocytes, indicating that they differentiate rapidly upon entry to the parenchyma of infected lungs; in contrast, cells that differentiate from Ly6C^hi^ monocytes turn over more slowly, but diminish in frequency in less than one week. New cells (identified by pulse-chase labeling) acquire bacteria within 1-3 days of appearance in the lungs, indicating that bacteria regularly encounter new cellular niches, even during the chronic stage of infection. Our findings that mononuclear phagocyte populations at the site of *M. tuberculosis* infection are highly dynamic provide support for specific approaches for host-directed therapies directed at monocytes, including trained immunity, as potential interventions in TB, by replacing cells with limited antimycobacterial capabilities with newly-recruited cells better able to restrict and kill *M. tuberculosis*.

**Author summary:** During certain chronic infections such as tuberculosis, inflammatory cells, including macrophages and dendritic cells, are recruited to infected tissues where they aggregate to form tissue lesions known as granulomas. Although granulomas can persist long term, the dynamics of the cell populations that comprise granulomas are not well understood. We used a combination of methods to discover that, during chronic infection of mice with *Mycobacterium tuberculosis*, the monocyte, macrophage, and dendritic cell populations are highly dynamic: recently-proliferated cells traffic rapidly to infected lung tissues, yet they persist with a half-life of less than one week. We also found that recently-proliferated cells become infected with *M. tuberculosis* as soon as one day after their arrival in the lungs, indicating that the bacteria are regularly moving to new cellular niches, even during the chronic stage of infection. The dynamic nature of the cell populations that encounter *M. tuberculosis* suggests that interventions such as trained immunity have potential therapeutic roles, by replacing cells that have poor antimycobacterial activity with cells with enhanced antimycobacterial activity. These interventions could improve the outcomes of treatment of drug resistant tuberculosis.

## Introduction

Mononuclear phagocytes (MNP) harbor *Mycobacterium tuberculosis* in tissues of humans [1] and experimental animals [2-4]; and MNP are essential elements of granulomas, the characteristic tissue lesions in tuberculosis [5, 6]. Although macrophages have been characterized as prominent cellular hosts for *M. tuberculosis in vivo*, recent studies have revealed the roles of distinct populations of MNP, including tissue-resident (i.e., lung alveolar) macrophages, monocyte-derived recruited macrophages, and dendritic cells, as host cells for the bacteria in experimental animals [4] and humans [7]. Although cells in these subsets exhibit functional differences during *M. tuberculosis* infection, including the ability to transport bacteria from the lungs to the local lymph nodes [8-10] and their ability to present antigens for activation of CD4 T cells [11], there is little known regarding the population dynamics of MNP in tuberculosis or any other chronic infection.

Recent studies of blood monocytes that emigrate from the bone marrow during homeostasis have revealed the potential for these cells to differentiate from Ly6C^hi^ monocytes to several distinct subsets of intravascular and tissue parenchymal cells. A proportion of Ly6C^hi^ monocytes differentiate into Ly6C^lo^ monocytes, which remain in the blood and vascular space of peripheral tissues, where they are considered to ‘patrol’ the vascular space and respond to inflammatory stimuli [12]. In addition, Ly6C^hi^ monocytes emigrate from the vascular space during homeostasis and differentiate into lung macrophages and dendritic cells [13]. *M. tuberculosis* infection markedly increases accumulation of recruited macrophages and dendritic cells in the lungs [2, 4, 9, 14, 15], but it is unclear whether the recruited cells are long-lived, or whether they require continuous replenishment by recruitment, local proliferation, or both. Since *M. tuberculosis* infection is accompanied by apoptosis [16], necrosis [17], and egress from the lungs to the local lymph node [8-10], we hypothesized that mononuclear cell populations in the lungs are dynamic, and their abundance and differentiation may contribute to the outcomes of infection.

Initial evidence that MNP populations can be dynamic at a tissue site was obtained in studies by Dannenberg, who used intravenous injection of tritiated thymidine to track the recruitment and decay of MNP at sites of *M. bovis* BCG injection in the skin of rabbits [18]. Although these experiments revealed evidence for MNP turnover, they could not reveal the differentiation of the labeled cells, and the decay of labeled cells at the site of BCG injection may have been at least partly due to clearing of the attenuated bacteria used as the stimulus.

Here, we studied monocyte trafficking, differentiation, and turnover in multiple compartments: blood, the lung vasculature, lung parenchyma, lung-draining lymph nodes, and distant lymph nodes, in uninfected and *M. tuberculosis*-infected mice, using adoptive transfer and *in vivo* pulse-chase labeling, coupled with flow cytometry and immunostaining of tissue sections. We selected time points that followed the rapid accumulation of cells that occurs during the early response to infection and that followed the development of adaptive immune responses. Our results extend previously-published results [15] by revealing distinct characteristics of endovascular and tissue parenchymal cells, by revealing that the MNP population is actively turning over during the chronic phase of *M. tuberculosis* infection, and by finding that new MNP rapidly acquire bacteria at the site of infection. The findings that MNP populations are dynamic during the chronic stage of infection suggests that mechanisms that disrupt MNP trafficking or differentiation may underlie reactivation and progression of TB, and also indicate the potential to employ trained immunity [19] as a therapeutic intervention in tuberculosis, by replacing cells that support intracellular bacterial growth with new cells that are more capable of restricting bacterial growth.

## Results

### Adoptively transferred monocytes differentiate into diverse mononuclear cells in *M. tuberculosis*-infected mice

To characterize the patterns and kinetics of trafficking and differentiation of monocyte-derived cells during the chronic phase of infection in a non-lymphoid tissue (the lungs), we purified Ly6C^hi^ and Ly6C^lo^ monocytes from the blood and bone marrow (BM) of CD45.1^+^ mice and transferred them intravenously (i.v.) to CD45.2^+^ recipients either 4 or 8 weeks after aerosol infection with *M. tuberculosis* H37Rv (Fig S1A). Mononuclear cells were ∼92-99% Ly6C^hi^ monocytes; the remainder were Ly6C^lo^ monocytes, dendritic cells (DC) and cells expressing markers of stem cell progenitors. We then characterized the adoptively-transferred CD45.1^+^ cells as they trafficked and differentiated in five compartments: peripheral blood, the lung vasculature, the lung parenchyma, the lung-draining mediastinal lymph nodes (MLN) and pooled peripheral lymph nodes that do not drain the lungs (pLN) (Figs S1 and S2A). Cells within the lung parenchyma were distinguished from intravascular cells by the i.v. administration of anti-CD45 antibody immediately prior to euthanasia.

At the time of the earliest sampling (∼40 h following transfer), the transferred cells in the blood and lung vasculature resembled the population of transferred cells: most were Ly6C^hi^ monocytes (Ly6C^hi^ CD11b^+^ CD11c^−^ MHCII^−^); the remainder were Ly6C^lo^ monocytes (Ly6C^lo^ CD11b^+^ CD11c^lo^ MHCII^−^), while no other monocyte-derived cell subset was detectable in the blood or the lung vascular space (Figs 1A, 1B, and S2A).

**Fig 1.**
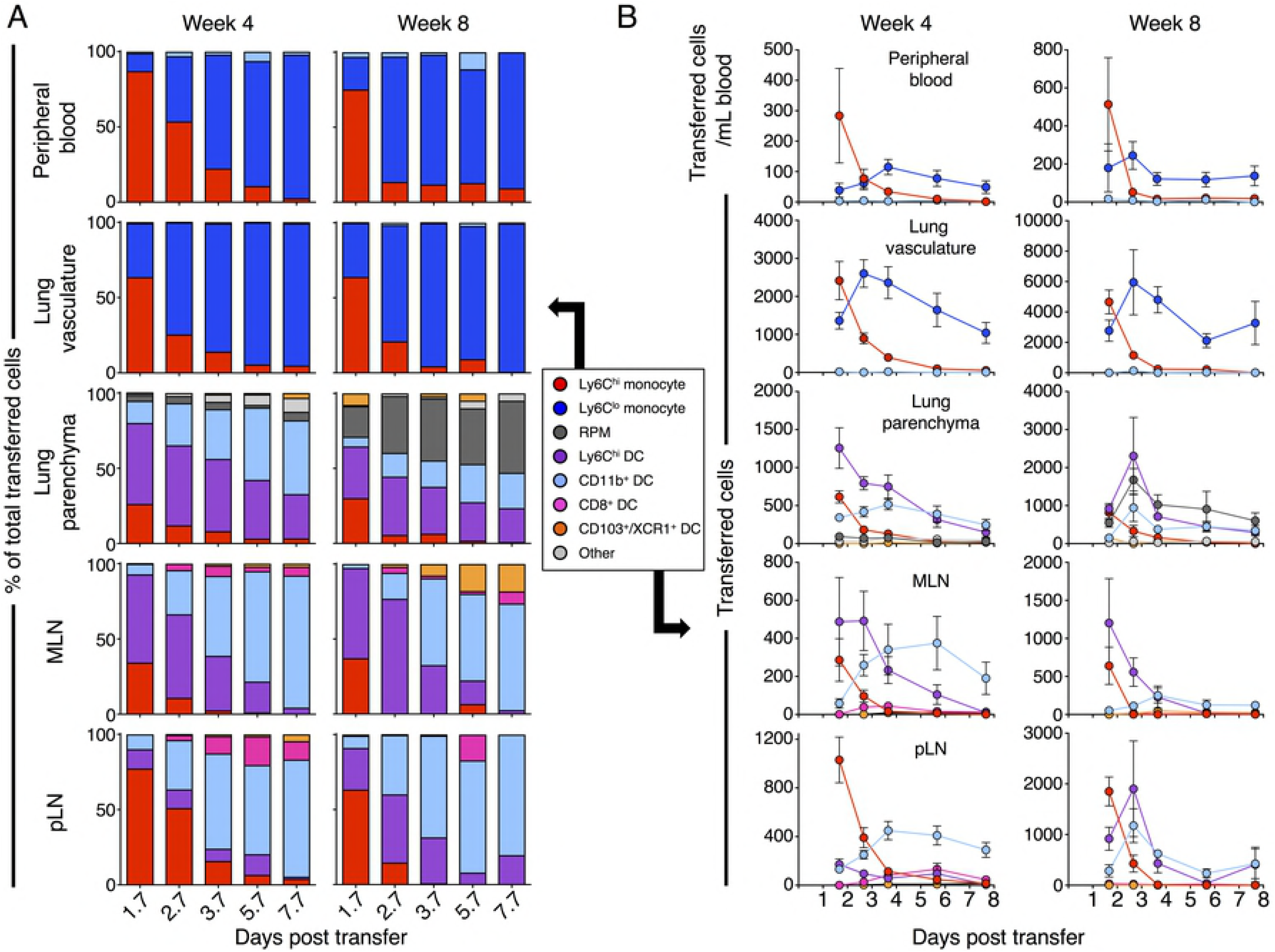
Adoptively transferred bone marrow monocytes differentiate into diverse cell populations in tissues of *M.tuberculosis*-infected mice. Bone marrow monocytes from CD45.1^+^ mice were depleted of stem cells and transferred to *M. tuberculosis*-infected CD45.2^+^ mice, 4 or 8 weeks after infection. CD45.1^+^ cells were identified by flow cytometry at multiple time points after transfer; cells in the lung vasculature were distinguished from cells in the lung parenchyma by the presence of labeling with a pan-CD45 antibody injected intravenously immediately before euthanasia. A) Subset frequency, as the proportion of the total CD45.1^+^ cells recovered; B) total subset numbers of CD45.1^+^ donor cells in blood, lung vasculature, lung parenchyma, lung-draining mediastinal lymph node (MLN) and total pooled peripheral LN (pLN). Data are from single experiments with 5 mice per time point. Proportion of transferred cells are means, and absolute cell numbers are means ± SEM.

The composition of transferred mononuclear cell populations in the lung parenchyma and MLN were similar, but notably different between weeks 4 and 8 of *M. tuberculosis* infection. At the first time point after transfer into mice infected for 4 weeks, >50% of the transferred cells in these tissues had differentiated into Ly6C^hi^ DCs (Ly6C^hi^ CD11b^+^ CD11c^+^ MHCII^hi^); the remainder of the transferred cells in these tissues were predominantly Ly6C^hi^ monocytes, while a small fraction had differentiated into CD11b^+^ DC (Ly6C^lo^ CD11b^+^ CD11c^+^ MHCII^hi^) in both tissues or into Ly6C^lo^ CD11b^+^ CD11c^lo^ MHCII^lo^ recruited parenchymal macrophages (RPM) in the lungs. Transferred cells differentiated almost identically in the MLN when transferred at 4 and 8 weeks post infection. However, at 8 weeks post *M. tuberculosis* infection, both Ly6C^hi^ DC and CD11b^+^ DC fractions were diminished in the lung parenchyma while the portion of transferred monocytes that became RPM was substantially increased.

While both Ly6C^lo^ monocytes and RPM express the phenotypic markers previously used to define lung recruited macrophages [4], they differ in their location within the lung vasculature or parenchyma. In the steady state, ∼10% of the Ly6C^lo^ monocytes/RPM are within the lung parenchyma (Fig S2B). However, as disease progresses, an increasing proportion of these cells are in the parenchyma, which is further evidenced by the increased number and proportion of donor-derived RPM during the later phase of infection (Fig 1A).

Recent publications have characterized heterogeneity within both the Ly6C^hi^ and Ly6C^lo^ monocyte populations in the bone marrow and circulation. Ly6C^lo^ monocytes are believed to be derived from Ly6C^hi^ monocytes, and in the steady state, the majority of the Ly6C^lo^ monocytes differentiate into I-Ab^−^ vascular-patrolling monocytes in an NR4A1-dependent manner [20, 21]. Ly6C^hi^ monocytes, on the other hand, have been shown to differentiate into monocyte-derived DCs and tissues macrophages in an NR4A1- and Flt3L-independent manner, with high expression of the transcription factor PU.1 predisposing these cells to differentiate into DC [22]. Intranuclear staining for the PU.1 revealed that DC populations in the lung parenchyma expressed the highest levels of PU.1 amongst mononuclear cells, while RPM expressed the lowest levels of PU.1 amongst mononuclear phagocytes (Fig S2C). These data support the assertion that Ly6C^hi^ monocytes contribute to bona fide DC and RPM cell subsets in the lungs of *M. tuberculosis*-infected mice.

In contrast to the MLN, Ly6C^hi^ monocytes were the dominant transferred cell subset in the pLN week 4 after infection, and only a small fraction of the cells were Ly6C^hi^ DC or CD11b^+^ DC. However, by week 8 of *M. tuberculosis* infection, transferred cells in the pLN differentiated into Ly6C^hi^ DC, suggesting that the inflammatory environment within pLN is distinct from MLN 4 weeks post infection, but the inflammatory milieu is more similar by week 8 of infection. The presence of *M. tuberculosis* in the pLN (data not shown) has been previously reported [10] and the inflammation from infection and the effector cytokines produced by *M. tuberculosis*-reactive T cells may contribute to the differentiation of Ly6C^hi^ DC in the LN. These results indicate that circulating bone marrow-derived monocytes enter an infected peripheral tissue and differentiate rapidly, and that differentiation into DC or RPM happens during or after egress from the vascular space in the lungs of *M. tuberculosis*-infected mice.

At later time points after transfer during both phases of *M. tuberculosis* infection, donor cells underwent further differentiation in tissue-dependent patterns, followed by gradual declines in their abundance in each compartment. In the peripheral blood and lung vascular space, Ly6C^hi^ monocytes progressively became less frequent as a fraction of the recovered cells, with nearly all of the remaining transferred cells present as Ly6C^lo^ monocytes 6-8 days after transfer (Fig 1A). In both of these compartments, the number of Ly6C^lo^ monocytes exhibited an inverse pattern relative to that of Ly6C^hi^ monocytes, with Ly6C^lo^ cells reaching a peak by days 3-4 followed by a gradual decline to approximately 30% of the peak level by day 8. This pattern is consistent with differentiation of Ly6C^hi^ to Ly6C^lo^ monocytes in the blood, as previously reported [12].

Over time, Ly6C^hi^ and CD11b^+^ DC exhibited an inverse relationship in the lung parenchyma and MLN. The total number of Ly6C^hi^ DC and their proportion of transferred cells continuously ebbed over time while CD11b^+^ DC numbers increased until day 4 before diminishing in number, while also constituting the majority of transferred cells. Surprisingly, although the kinetics of Ly6C^hi^ monocytes and CD11b^+^ DC turnover were similar in the pLN and MLN, only a small fraction of transferred cells differentiated into Ly6C^hi^ DC in the pLN.

The pattern and kinetics of adoptively transferred monocyte trafficking and differentiation in the MLN were similar at later phases of infection (8 weeks post infection), though, in the lung parenchyma and in pLN there was a delay in the peak number of Ly6C^hi^ DC to nearly 3 days after transfer (Fig 1). Transferred monocytes also differentiated into CD11b^+^ DC more rapidly during this later phase of infection, peaking at day 3 instead of 4 days after transfer. The most striking difference between the two infectious phases, however, was the greater increase in RPM in the lung parenchyma at the later phase of infection. This suggests the inflammatory environment within the lung is changing throughout infection, leading to an evolving population of mononuclear cells with different phenotypes.

We determined the half-life of the Ly6C^hi^ monocytes as well as the total transferred CD45.1^+^ mononuclear cell population in each of the 5 compartments examined by fitting a first order exponential decay function to the data on the cell numbers identified by flow cytometry. Although there was substantial variation in the half-lives of the CD45.1^+^ cells in each compartment, the data were best described by a best-fit line for all the compartments combined, which yielded a consensus half-life of 112.18 hours (Fig S2C). In contrast, the decay of Ly6C^hi^ monocytes (by differentiation, egress, or other mechanisms) was most rapid in the pLN and was similar to that in the lung vasculature and parenchyma. By comparison, decay of Ly6C^hi^ monocytes was slower in the MLN and blood. When the qualities of the best-fit equations were compared, it was determined that a single best-fit equation for all the Ly6C^hi^ monocytes in all tissues was a better fit than equations for each tissue. This best-fit equation for all the Ly6C^hi^ monocyte data yielded a half-life of 15.31 hours (Fig S2D). Together, these results indicate that Ly6C^hi^ monocytes have a half-life of less than one day, while cells that differentiate from Ly6C^hi^ monocytes have a half-life of approximately 5 days during the chronic stage of infection with *M. tuberculosis*.

The results of these monocyte adoptive transfer studies are consistent with studies of the homeostatic state in mice indicating that Ly6C^hi^ monocytes that egress the bone marrow can differentiate into Ly6C^lo^ monocytes that remain in the blood and vascular space for prolonged periods [12]. In addition, they reveal that during the chronic phase of *M. tuberculosis* infection, monocytes that enter the lung parenchyma and lung-draining MLN preferentially differentiate into Ly6C^hi^ DC, followed by CD11b^+^ DC. In contrast, monocytes that traffic to a lymph node distant from the lungs undergo more diverse pathways of differentiation, suggesting that the local tissue environment contributes to monocyte differentiation into distinct cell subsets.

### Recently-proliferated monocytes have similar turnover in lymphoid and peripheral tissues during *M. tuberculosis* infection

Our adoptive transfer experiments established the kinetics and potential for monocytes to traffic and differentiate in distinct compartments in the context of chronic infection with *M. tuberculosis* but could not provide insight into the quantitative contributions of monocytes compared with resident cells in the lungs or lymph nodes during infection. To further understand the dynamics of mononuclear cell trafficking, proliferation, and differentiation, we briefly pulsed uninfected and *M. tuberculosis*-infected mice at multiple stages of chronic infection with the nucleoside EdU (5-ethynyl-2’-deoxyuridine) and characterized monocytic cells with EdU-labeled DNA at multiple time points in the bone marrow, blood, lung vascular space, lung parenchyma, and MLN.

Quantitation of total Ly6C^hi^ cells in the bone marrow revealed no differences between uninfected mice and *M. tuberculosis*-infected mice, at any stage of chronic infection (up to 16 weeks) (Fig 2A). In contrast to the bone marrow, in the blood we found that chronic *M. tuberculosis* infection was associated with increases in Ly6C^hi^ monocytes, with the most marked increase observed at the latest time point of infection examined (16 weeks post infection) (Figs 2A and S3A). The finding in the blood is consistent with observations with human tuberculosis patients, in which peripheral blood monocytosis is associated with active tuberculosis and with the severity of tuberculosis in children [23]. The findings in the blood are also consistent with recent reports of increased monocyte/lymphocyte ratios in the blood of patients with active TB [24] or at high risk of progression to active TB [25].

**Fig 2.**
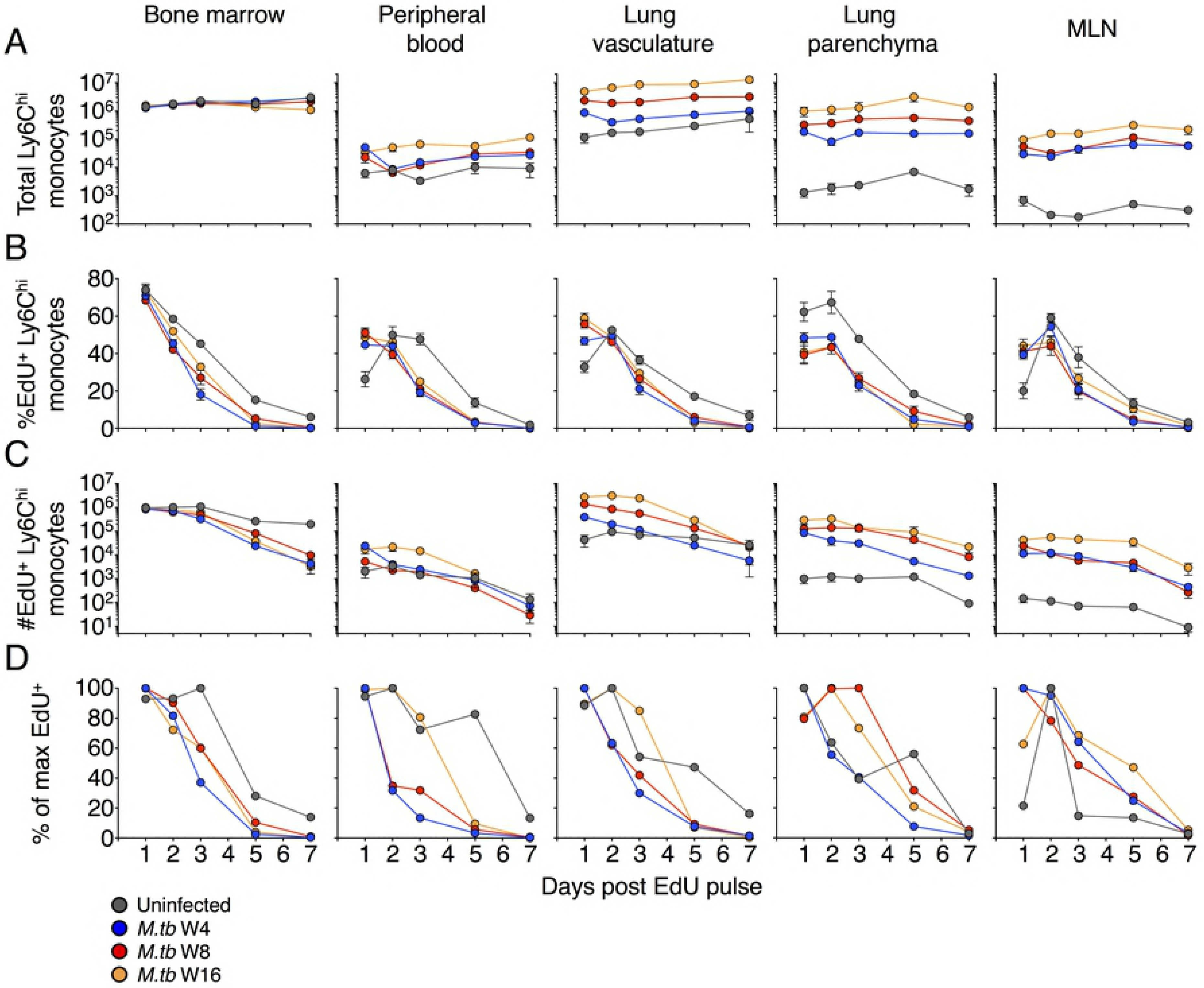
Turnover kinetics of Ly6C^hi^ monocytes in lymphoid and peripheral organs during *M. tuberculosis* infection. Mice were injected with EdU at multiple phases of *M. tuberculosis* infection and EdU incorporation by dividing monocytes was evaluated by flow cytometry at multiple time points after the EdU pulse. A) Total numbers (EdU^+^ and EdU^−^) of Ly6C^hi^ monocytes in distinct tissue compartments of uninfected mice and mice infected with *M. tuberculosis* for 4, 8, or 16 weeks prior to the EdU pulse. B) Kinetics of turnover of EdU^+^Ly6C^hi^ monocytes in distinct tissue compartments, as the % of Ly6C^hi^ monocytes that are EdU^+^; C) kinetics of turnover of EdU^+^ Ly6C^hi^ monocytes in distinct tissue compartments, as the number of EdU^+^ Ly6C^hi^ monocytes. Data are presented as means ± SEM from 1-4 experiments per infection phase with 4-5 mice per time point per experiment. D) Normalized kinetics of turnover of EdU^+^Ly6C^hi^ monocytes in distinct tissue compartments of uninfected mice or 4, 8, and 16 weeks after infection with *M. tuberculosis.* Variations in total cell numbers between time points were normalized by finding the maximum fraction of EdU^+^ Ly6C^hi^ monocytes within total live cells across all time points per phase and then presenting the means relative to the maximum % of live cells. Numbers of bone marrow monocytes are per leg, both tibia and fibula; and in peripheral blood as per mL.

The finding of increased numbers of blood monocytes during infection without an increase in the number of mature monocytes at their site of production in the bone marrow is consistent with a shorter dwell time of mature monocytes in the bone marrow, and/or a longer dwell time in the blood. Analysis of the frequency of labeled monocytes after a brief pulse of EdU revealed a significant increase in the rate of decay or egress of monocytes from the bone marrow associated with infection (Figs 2B and 2C). The half-life of Ly6C^hi^ monocytes in the BM decreased from 47.59 h in uninfected mice to 26.06 h at the most acute phase of infection examined, though, the half-life had increased to 34.27 h by the 16^th^ week of infection (Fig S3B and table S1A). We also determined the half-lives of Ly6C^hi^ monocytes by fitting the equation to the total numbers of EdU^+^ Ly6C^hi^ monocytes; this revealed a similar trend of increased monocyte turnover during *M. tuberculosis* infection (Fig S3C and Table S1B). These results suggest that acute and chronic *M. tuberculosis* infection are associated with increased production of monocytes in the bone marrow, which is balanced by increased egress from the bone marrow to the blood and to peripheral tissues. This interpretation was supported by analysis of the frequency of EdU-labeled monocytes in the blood. First, comparison of the frequency of EdU pulse-labeled monocytes in the blood of uninfected and infected mice revealed a lag before EdU-labeled monocytes peaked in the blood of uninfected mice; this lag was not evident in the blood of *M. tuberculosis*-infected mice, consistent with more rapid entry of monocytes into the blood from the bone marrow (Fig 2B and 2D). Second, the rate of decay of the frequency of EdU-labeled monocytes from the blood was greater in infected mice (ranging from 17.6-23.6 h) compared with that in uninfected mice (34.1 h) (Figs S3B and 2C, and S1 Table). Together, these results show that increased numbers of monocytes in the blood of *M. tuberculosis*-infected mice result from increased production of monocytes in the bone marrow, balanced by increased release of mature monocytes from the bone marrow into the blood, and not by prolonged retention of monocytes in the blood.

Ly6C^hi^ monocytes accumulated in the peripheral tissues during *M. tuberculosis* infection, with ∼30-fold increase in the lung vasculature over that in uninfected mice (S2 Table). The kinetics of EdU pulse-labeled monocytes in the lung vascular space closely mimicked those in the blood, including the delay to the peak in the frequency of labeled Ly6C^hi^ monocytes in uninfected mice that was absent in infected mice (Figs 2B and 2D), and the shorter half-life of monocytes in blood of infected mice.

In the lung parenchyma, *M. tuberculosis* infection was accompanied by a striking increase in the total number of monocytes, as previously reported [4, 15] the number of Ly6C^hi^ monocytes in the lung parenchyma progressively increased between 4 and 16 weeks of infection where they were more than 550 times more numerous than in uninfected mice (Fig 2A and S2 Table). The peak in the frequency of EdU pulse-labeled monocytes was lower in the lung parenchyma of infected mice compared with that of uninfected mice; this is most likely due to the larger population of monocytes in the lungs of infected mice into which the EdU^+^ cells are diluted. After peaking on day 2 post EdU pulse, the rate of decay of Ly6C^hi^ EdU^+^ cells was increased in *M. tuberculosis*-infected mice (Figs 2B-C, S3B-C and table S2).

The behavior of Ly6C^hi^ EdU^+^ monocytes in the lung-draining MLN closely mimicked those in the lung parenchyma: the number of cells in both tissues increased markedly and progressively with *M. tuberculosis* infection, and the kinetics of appearance and decay of Ly6C^hi^ EdU^+^ cells were similar. In both tissues, infection accelerated the rate of decay of Ly6C^hi^ cells, and by 7 days after the EdU pulse, the number of Ly6C^hi^ cells decreased 10 to 100-fold compared to the peak value reached at 2 days post-pulse. During both acute and chronic phases of *M. tuberculosis* infection, Ly6C^hi^ monocytes rapidly proliferate in the BM, and enter the periphery to differentiate into various cell subsets at an increased rate relative to uninfected mice.

### Monocyte-derived macrophage and dendritic cells accumulate in the lungs of *M. tuberculosis* infected mice but continuously turn over

Since the quantitative decay of Ly6C^hi^ EdU^+^ monocytes in tissues can be due to differentiation, egress, cell division, and/or death, we analyzed the differentiation of EdU^+^ cells over time in the lung vasculature and parenchyma of uninfected and *M. tuberculosis*-infected mice. Total numbers of Ly6C^lo^ monocytes in the peripheral blood were modestly increased only at the later chronic phase of *M. tuberculosis* infection that we examined (Fig S4A). In the lungs, however, the numbers of the Ly6C^lo^ CD11b^+^ CD11c^lo^ MHCII^lo^ subset we have previously characterized as recruited macrophages [4, 9, 11] and Ly6C^lo^ monocytes (CD45 IV^−^) continuously increased in the lungs of *M. tuberculosis*-infected mice as disease progressed (Fig 3A and S4A). As with our adoptive transfer data, the majority of these cells were in the lung vasculature but the proportion of these cells that were in the parenchyma as RPM increased from ∼10% in uninfected mice to ∼25% in mice infected with *M. tuberculosis* for 16 weeks (Fig S1B).

**Fig 3.**
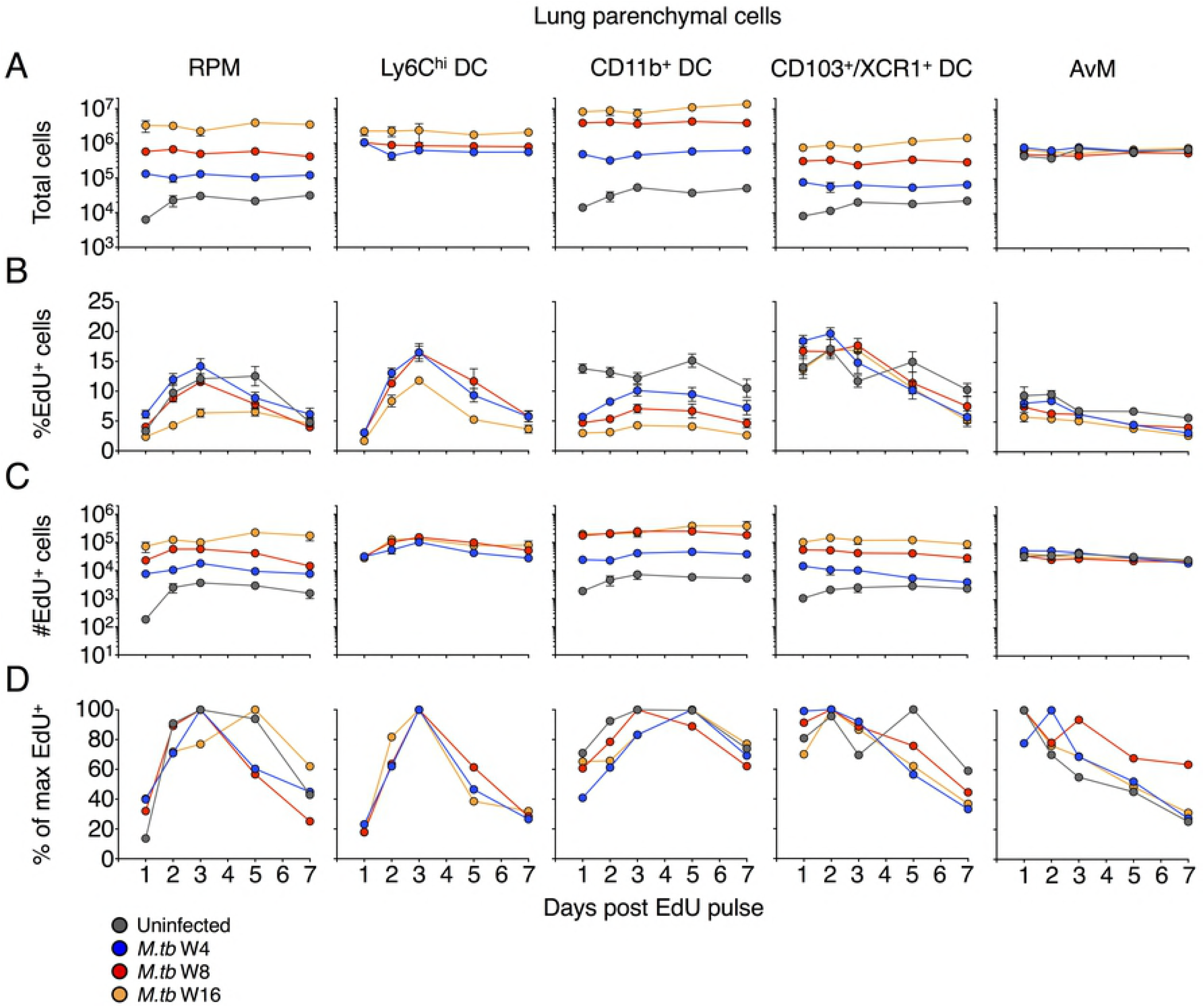
Kinetics of macrophage and dendritic cell turnover in the lung parenchyma of *M. tuberculosis*-infected mice. Mice received a pulse of EdU after 4, 8, or 16 weeks of *M. tuberculosis* infection, and EdU-labeled cells in distinct mononuclear cell subsets were evaluated by flow cytometry at multiple time points following the pulse of EdU. A) Total number of cells (EdU^+^ and EdU^−^) in the designated subsets in the lung parenchyma of uninfected and *M. tuberculosis*-infected mice, RPM: recruited parenchymal macrophages; AvM: alveolar macrophages. B) Frequency (%) of EdU^+^ cells in each subset; C) total numbers of EdU^+^ cells in each subset. Data are presented as means ± SEM from 1-4 experiments per infection phase with 4-5 mice per time point per experiment. D) Normalized kinetics of turnover of EdU^+^ cells in distinct subsets, as the mean percent of the maximum frequency of labeled cells in lungs of uninfected or *M. tuberculosis*-infected mice. Data were normalized as in Fig 2D.

By 24 hours after EdU pulse, only a small fraction of these Ly6C^lo^ monocyte and RPM subsets were EdU^+^, matching the delayed differentiation of Ly6C^hi^ monocytes to Ly6C^lo^ monocytes and RPM we observed in our adoptive transfer experiments (Figs 3B-D and S4B-D). The peak in the frequency of EdU^+^ RPM appeared later after the EdU pulse than did the peak of Ly6C^hi^ cells in the lung parenchyma (3 d vs 2 d, respectively), consistent with Ly6C^hi^ cells as precursors of at least a fraction of the RPM. Although the peak of EdU labeling was delayed in these cells, EdU^+^ RPM declined in frequency and total number, indicating that this cell population undergoes continuous turnover during both acute and chronic phases of infection. EdU^+^ RPM exhibited several differences later in infection (16 weeks) compared with the pattern 4-8 weeks post infection; the peak frequency of EdU^+^ RPM was delayed and significantly lower than in other phases of infection. At this time point, the total number of RPM and the number of EdU^+^ RPM was higher than observed 4 or 8 weeks post infection. Together, these results indicate that the dynamics of mononuclear cells at the site of infection change during the chronic phase of infection, resulting in a larger population of RPM that turns over more slowly and comprises a larger proportion of the parenchymal cells (Figs S4E-F).

Consistent with the data from adoptive transfer of bone marrow monocytes, the EdU pulse labeled up to 10-15% of the Ly6C^hi^ DC in the lung parenchyma with peak labeling observed 3 days after the EdU pulse (Fig 3B and 3C). Also consistent with the results of adoptive transfer, Ly6C^hi^ DC are transient, with the proportion of EdU^+^ cells decaying to ∼30% of the peak level one week after EdU pulse. Furthermore, these cells did not accumulate with disease progression as observed with other mononuclear cell populations (RPM, CD11b^+^ DC, CD103^+^/XCR1^+^ DC), displayed similar kinetics of EdU incorporation and decay between infection phases, and were undetectable in the lungs of uninfected mice. These cells likely represent a transient transition state from Ly6C^hi^ monocytes to CD11b^+^ DC or RPM and are only detectable during infection.

CD11b^+^ DC progressively increased in number within the lung parenchyma of *M. tuberculosis*infected mice and EdU^+^ CD11b^+^ DC reached a peak frequency ∼3 days after the EdU pulse (Fig 3B). A substantial fraction of CD11b^+^ DC were EdU^+^ within 24 h of EdU administration, suggesting that these cells can proliferate locally in the lungs, as previously reported [26] or have differentiated rapidly from recently-proliferated precursors. This fraction was diminished during infection, and a smaller portion of CD11b^+^ DC had recently proliferated, especially at later phases of infection, reflecting a greater role for monocyte recruitment and differentiation to DC during *M. tuberculosis* infection.

We examined two additional subsets of mononuclear phagocytes in the lung parenchyma: CD103^+^/XCR1^+^ DC and alveolar macrophages (AvM). For some of the experiments we utilized antibodies specific for XCR1 instead of CD103 as XCR1^+^ DC have been shown to largely consist of CD103^+^ CD11b^−^ DC in the LN and peripheral organs [27, 28]. Up to 12-15% of the CD103^+^/XCR1^+^ DC in the lung parenchyma were labeled with an EdU pulse; unlike the other cell subsets studied, the frequency of EdU^+^ CD103^+^/XCR1^+^ DC did not differ in uninfected compared with infected mice, although the total number of CD103^+^ DC and the number of EdU^+^ CD103^+^/XCR1^+^ DC increased in a time-dependent manner with *M. tuberculosis* infection. Notably, the peak of EdU-labeled CD103^+^/XCR1^+^ DC did not exhibit the lag observed with RPM, Ly6C^hi^ DC, or CD11b^+^ DC (Fig 3), suggesting that this subset of cells in the lung parenchyma does not depend on differentiation from a cell with another phenotype, and that CD103^+^/XCR1^+^ DC proliferate locally in the lungs, even in the absence of infection, as previously reported [26]. Compared with the other cell subsets examined, a smaller fraction (<10%) of AvM incorporated EdU, and the frequency and number of EdU^+^ AvM was unaffected by *M. tuberculosis* infection at any time point examined. Also, unlike the other cell subsets examined, there was no detectable increase in the total number of AvM in lungs of infected mice compared with uninfected mice (Fig 3). The majority of mononuclear cell subsets in the lung parenchyma of infected mice had low (1-2%) expression of Ki67, while ∼10% of CD103^+^/XCR1^+^ DC were Ki67^+^ (Fig S4G). Thus, the majority of professional phagocyte subsets we identified were derived by differentiation from Ly6C^hi^ monocytes, while CD103^+^/XCR1^+^ DC and AvM proliferated in situ with minimal contribution from monocytes.

Finally, we calculated the total number of potential monocyte precursors within the lung vasculature and compared it to the total number of putative monocyte-derived populations in the parenchyma. We determined the area-under-the-curve for lung vasculature Ly6C^hi^ and Ly6C^lo^ monocytes as well as DCs throughout 7 days post EdU pulse for each phase and compared it to the area-under-the-curve for parenchymal monocytes, RPM, Ly6C^hi^ DC, CD11b^+^ DC and CD103^+^/XCR1^+^ DC (Fig S4H). The turnover of circulating EdU^+^ precursors of vascular monocytes, MP and DC populations was greater than the combined number of these parenchymal populations in uninfected mice and at all phases of *M. tuberculosis* infection. Thus, the turnover of recently-proliferated vascular monocytes is sufficient to account for the turnover in monocyte-derived cells in the lung parenchyma.

### Maturation and activation of monocytes as they differentiate into DC and MP subsets

We further characterized the differentiation of recently-proliferated monocytes and their descendants into cells with antigen-presenting potential by comparing the expression of MHCII in new (EdU^+^) cells relative to all cells of each subset. Ly6C^hi^ monocytes within in the lungs increased their expression of MHCII as they moved from the vasculature into the parenchyma and differentiated into Ly6C^hi^ and CD11b^+^ DC (Fig 4). At 12h after an EdU pulse, in the lung parenchyma and vasculature EdU^+^ Ly6C^hi^ monocytes expressed significantly less surface MHCII than the total population of Ly6C^hi^ monocytes. New Ly6C^hi^ DC, CD11b^+^ DC and CD103^+^ DC also had significantly lower expression of MHCII when compared to total MHCII expression of these subsets. However, by 24h all populations of new EdU^+^ had upregulated MHCII to the same level as their respective total population. These data support previous work demonstrating the upregulation of MHCII by Ly6C^hi^ monocytes as they transition from the vasculature into peripheral tissues [29].

**Fig 4.**
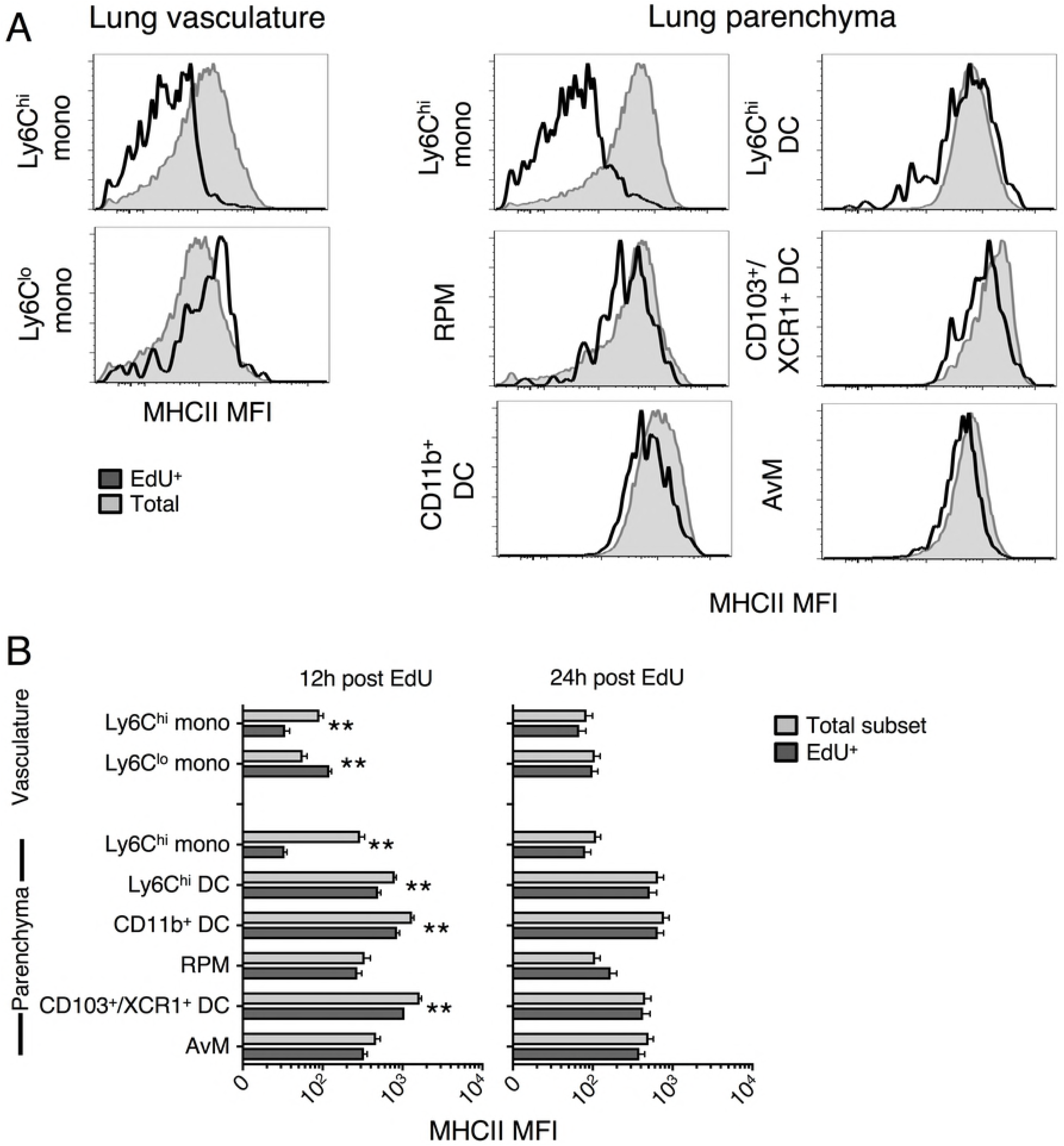
Recently-proliferated monocytes upregulate MHCII expression as they differentiate in the lungs. The relative expression of surface MHCII in EdU^+^ and total (EdU^+^ and EdU^−^) cells in multiple mononuclear cell subsets isolated from the lungs was determined by flow cytometry. A) Representative histogram of surface MHCII expression on cells in total lung mononuclear cell subsets and EdU^+^ cells in each subset, 4 weeks after infection with *M. tuberculosis* and 12 h after EdU pulse. B) Surface expression of MHCII (as mean fluorescence intensity (MFI)) on total and EdU^+^ cells in distinct mononuclear cell subsets, 4 weeks after infection with *M. tuberculosis* and 12 or 24 h after EdU pulse. Data are presented as mean ± SEM from 1-4 experiments with 4-5 mice per time point per experiment.

### Diversity of mononuclear cell subset kinetics in lung-draining lymph nodes

The lung-draining lymph nodes play pivotal roles in TB immunity. Live *M. tuberculosis* bacteria are transported from the lungs to the local lymph nodes by DC [10, 30], where infected migratory DC transfer antigens to uninfected resident DC to initiate T cell priming [8, 9]. The *M. tuberculosis* population within the MLN also represents a substantial bacterial burden [10, 31] and in humans may act as reservoirs of the pathogen for activation of latent TB [32]. Priming and development of CD4 T cell responses in the MLN is key to the eventual control of *M. tuberculosis* replication within the lungs, but also necessary for generating the proinflammatory adaptive immune responses that facilitate active TB disease and transmission. Understanding the relationships of antigen presenting cells within the MLN with those in the lungs may be essential for optimal development of effective vaccines or pharmaceutical treatments of TB.

We identified multiple mononuclear cell subsets in the MLN following *M. tuberculosis* infection and characterized their incorporation of EdU over time (Fig S5A). *M. tuberculosis* infection causes large increases in the population size of several mononuclear cell subsets in the MLN (Fig 5A) in addition to substantial increases of B and T lymphocyte populations (data not shown). The marked expansion of mononuclear cells affects at least 6 distinct subsets, develops by 4 weeks post-infection, and persists for at least 16 weeks (Fig 5A).

**Fig 5.**
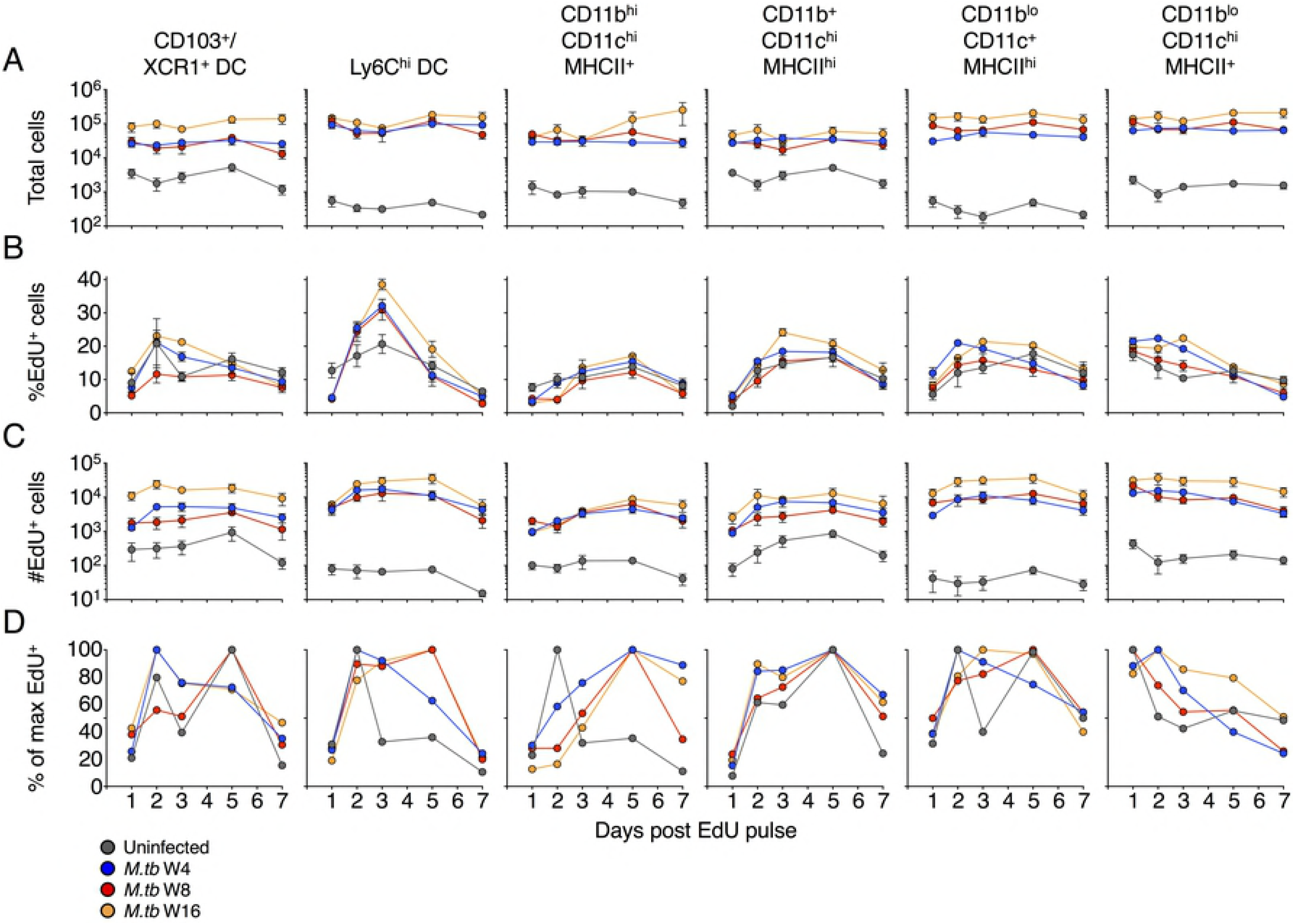
Kinetics of mononuclear cell subset turnover in the lung-draining mediastinal lymph node (MLN). Mice received a pulse of EdU after 4, 8, or 16 weeks of *M. tuberculosis* infection, and EdU-labeled cells in distinct mononuclear cell subsets were evaluated by flow cytometry at multiple time points following the pulse of EdU. A) Total number of cells (EdU^+^ and EdU^−^) in the designated subsets in MLN of uninfected and *M. tuberculosis*-infected mice. B) Frequency (%) of EdU^+^ cells in each subset; C) total numbers of EdU^+^ cells in each subset. Data are presented as means ± SEM from 1-4 experiments per infection phase with 4-5 mice per time point per experiment. D) Normalized kinetics of turnover of EdU^+^ cells in distinct subsets, as the mean percent of the maximum frequency of labeled cells in MLN of uninfected or *M. tuberculosis*-infected mice. Data were normalized as in Fig 2D.

Amongst all myeloid cells in the MLN that contain bacteria, Ly6C^−^ CD11b^hi^ CD11c^hi^ MHCII^+^ cells constitute the largest population infected with *M. tuberculosis* at all time points examined (Fig S5B), in agreement with previous studies that were limited to early time points [4, 9]. There were also minor populations of Ly6C^hi^ DC (CD11b^+^ CD11c^+^ MHCII^+^) infected with *M. tuberculosis*, while few of the CD11b^−^ DC (Ly6C^−^ CD11b^lo^ CD11c^+^ MHCII^hi^ and Ly6C^−^ CD11b^lo^ CD11c^hi^ MHCII^+^) [9] contained bacteria as detected by flow cytometry.

Ly6C^−^ CD11b^lo^ CD11c^hi^ MHCII^+^ DC rapidly incorporated EdU following the pulse (Fig 5), suggesting they proliferated within the MLN and supporting their classification as resident DC. The other mononuclear cell subsets examined all showed increasing EdU incorporation over time, albeit with different kinetics between populations and infection phases. Furthermore, each of these populations had different levels of EdU incorporation 24 h post pulse, potentially reflecting varied contributions of proliferation within the MLN. Of the mononuclear cell populations subsets examined, Ly6C^hi^ DC decayed most rapidly, consistent with their differentiation into other DC subsets. Other mononuclear subsets also decayed in the MLN during the 7 days following the EdU pulse, indicating that their populations remain dynamic as late as 16 weeks post-infection.

Mononuclear cells present in the MLN during *M. tuberculosis* infection might arrive from the blood via high endothelial venules (HEV) or from the lungs through afferent lymphatics [33, 34]. Monocytes expressing CD62L bind to heavily glycosylated molecules expressed on HEV and enter into lymph nodes from the blood [29, 35]. To determine the contribution of blood-derived, HEV-recruited monocytes to the mononuclear cell subsets present during *M. tuberculosis* infection we blocked CD62L in conjunction with an EdU pulse to identify recently-proliferated and/or recruited cells (Fig S5C).

As expected, there were profound reductions in total cell numbers in the MLN and peripheral LNs following anti-CD62L treatment (data not shown). However, there were no significant differences in the total numbers of Ly6C^hi^ monocytes in any of the compartments we evaluated (Fig S5D). To compare the relative contribution of HEV-migration on monocytes and mononuclear cell subsets within the LN that had drastically different total cell numbers we evaluated the relative fraction of EdU incorporation between anti-CD62L-treated and control mice. There was a small but significant reduction in the frequency of EdU^+^ Ly6C^hi^ monocytes in the MLN following CD62L blockade while there were no differences in the BM, blood, pLN or lungs (Fig S5E). In contrast to the small reduction in proportion of new Ly6C^hi^ monocytes in the MLN, there were no differences in EdU incorporation in other monocytes, RPM, CD11b^+^ DC in the lungs or in the multiple mononuclear cell subsets in the lymph nodes (data not shown). Thus, a small fraction of Ly6C^hi^ monocytes are directly entering into the MLN, while the majority of these cells arrive from the lungs through the afferent lymphatics. The vast majority of potential *M. tuberculosis*-harboring mononuclear subsets within the MLN are derived from cells that have passed through the lung lymphatics or proliferated within the MLN.

### Recently proliferated myeloid cells enter granulomas and become infected shortly after arrival

Since only a small minority of the cells that accumulate at the site of *M. tuberculosis* infection in the lungs harbor bacteria [4], we investigated whether the bacteria remain in the same cells for long periods, or whether they are acquired by new cells. We took advantage of EdU labeling to identify new cells and to localize and quantitate the presence of fluorescent protein-expressing bacteria in the recently-proliferated cells.

Immunofluorescence microscopy of lung sections revealed that by day three following the EdU pulse recently-proliferated (EdU^+^) CD11b^+^ professional phagocytes are evident within granulomas containing GFP-expressing *M. tuberculosis*, 4 weeks (Fig 6A) or 8 weeks (not shown) after infection. This timing corresponds to the peak numbers of EdU^+^ CD11b^+^ cells in the lung parenchyma which were comprised of both mononuclear cell subsets and neutrophils by flow cytometry (Figs 3, S4, and S6A). Consistent with the evidence from flow cytometry studies (Fig 3B), EdU^+^ cells became less abundant in lung lesions with increasing time after the EdU pulse (data not shown). As *M. tuberculosis* infection progressed, CD11b^+^ DC increasingly became the dominant cell type containing bacteria in the lung parenchyma, particularly at the expense of Ly6C^hi^ DC and to a lesser extent, neutrophils (Fig S6B). Recently-proliferated CD11b^+^ cells were also found to contain GFP-expressing *M. tuberculosis* in the lung-draining MLN 4 (Fig 5B) and 8 weeks post infection (Data not shown). Although EdU^+^ nuclei within CD11b^+^ lesions are in close proximity to GFP^+^ bacilli (Figs 5A and 5B), the indistinct cell boundaries in the lung sections precluded quantitation of new, EdU-labeled CD11b^+^ cells that have acquired bacteria.

**Fig 6.**
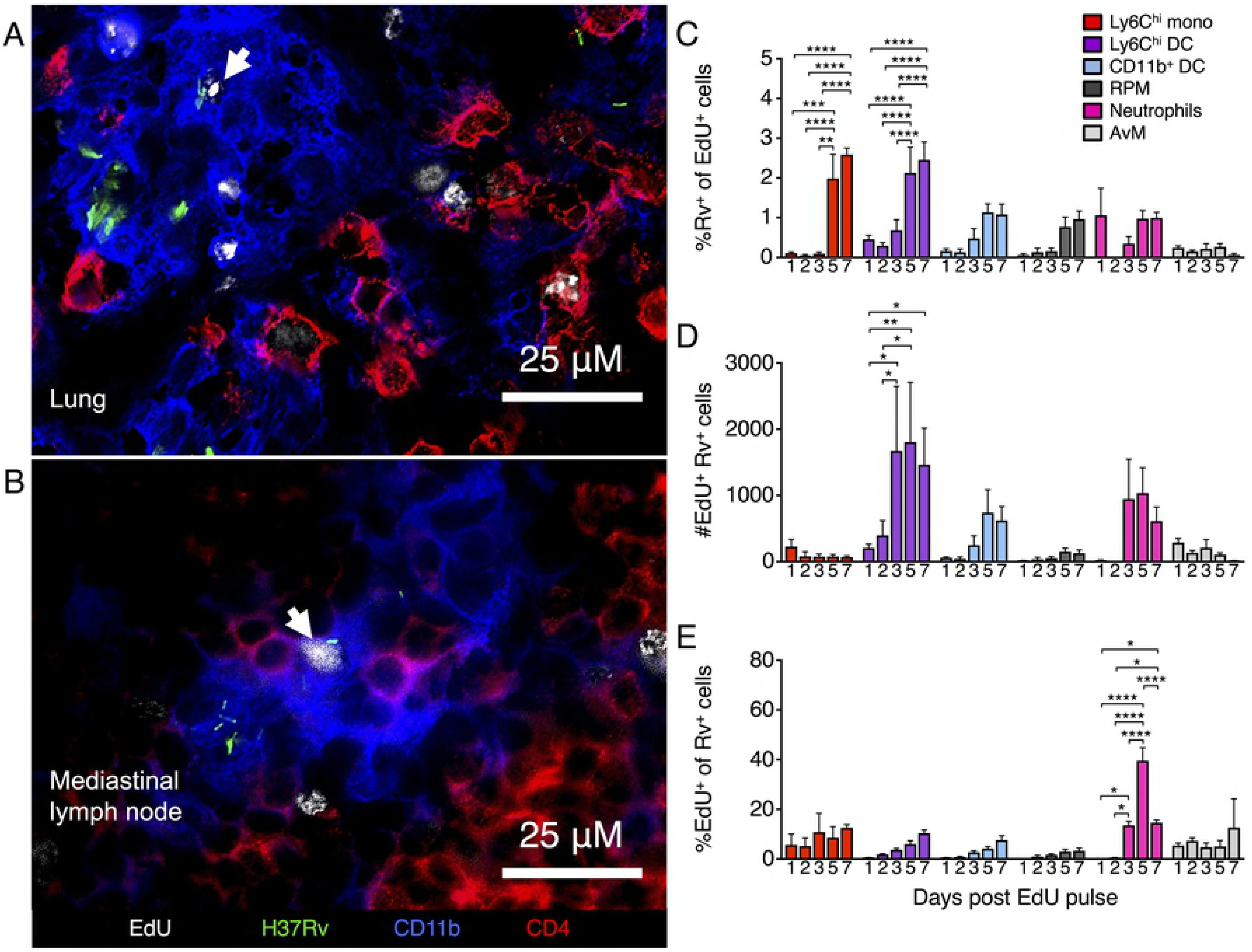
Recently proliferated mononuclear cells enter lung granulomas and become infected by *M. tuberculosis*. Mice infected with fluorescent protein-expressing *M. tuberculosis* were injected with EdU and its incorporation by dividing mononuclear cells evaluated by fluorescence microscopy or flow cytometry. Representative immunofluorescent staining of A) lung granulomas or B) CD11b^+^ lesions in the MLN of mice infected with GFP-expressing *M. tuberculosis*, 3 or 2 days after EdU pulse, respectively. Arrows indicate EdU^+^ nuclei adjacent to GFP-expressing *M. tuberculosis* bacilli. Images are representative of 1-2 mice per time point. C) Flow cytometric analysis of the frequency of EdU^+^ phagocyte subsets that were infected with Rv-mCherry at multiple time points following EdU pulse 4 weeks post infection. D) Flow cytometric analysis of total number of Rv^+^ EdU^+^ phagocyte subsets at multiple time points following EdU pulse 4 weeks post infection. E) Flow cytometric analysis of the frequency of Rv-mCherry-infected professional phagocytes that were EdU^+^ at multiple time points following EdU pulse 4 weeks post infection. Data are presented as means ± SEM of 4-5 mice per time point from a single infection for each phase. Statistics are 2-way ANOVA with Sidak’s multiple comparisons tests. *p<0.05, **p<0.01, ***p<0.001, ****p<0.0001.

To quantitate new (EdU pulse-labeled) cells that become infected soon after they appear in the lungs, we used flow cytometry to detect the coincidence of EdU labeling and *M. tuberculosis* H37Rv-mCherry in the cell subsets of interest. Because the numbers of EdU^+^ Rv^+^ cells in the the MLN were so infrequent, we were only able to quantify newly infected cells within the lung. Examination at either 4 weeks or 16 weeks post infection revealed that a fraction of the EdU^+^ Ly6C^hi^ monocytes, Ly6C^hi^ DC, CD11b^+^ DC, RPM, neutrophils and AvM contained bacteria as early as one day after the EdU pulse (Figs 5C-D and S6C-D). At both time points post infection, the frequency of infected EdU^+^ cells increased progressively in each of these cell subsets, up to 7 days post EdU (Figs 5C and S5C).

We also characterized the dynamics of infection with an alternative approach, by determining the fraction of bacteria-containing cell subsets that were comprised of EdU^+^ new cells. Most infected cell subsets contained increasing amounts of EdU^+^ cells over the week post EdU pulse, with up to 10-20% recently proliferated cells at both 4 and 16 weeks after infection (Figs 5E and S6E). In contrast, *M. tuberculosis*-infected Ly6C^hi^ monocytes week 16 post infection and neutrophils at both infection phases had a substantially higher proportion of new cells, up to 50% and 40% EdU^+^ respectively. Instead of continuously increasing, the proportion of these infected cells peaked in EdU staining before rapidly diminishing to 20-25% of the peak the following day. The large fraction of these particular subsets that is composed of new cells and the drastic loss of EdU staining is indicative of their relative short half-lives within granulomas before differentiating, migrating or dying.

These data demonstrate that newly-proliferated and -differentiated mononuclear cells are constantly trafficking to the infected lung where they are able to enter granulomas during the early and the later phase of infection. Importantly, these results also reveal that new mononuclear cells become infected shortly after their appearance at the site of infection, indicating that while the overall bacterial population remains at a steady state, the bacteria themselves are regularly entering new cellular environments. Since these data are only able to reveal the acquisition of bacteria by the new/EdU^+^ cells which are a small fraction of the total population at a given time, our results likely underestimate the frequency of bacterial transfer from cell to cell that is ongoing at the site of infection, even after development of adaptive immune responses.

## Discussion

In this work, we advanced the understanding of host cell and bacterial dynamics during the chronic stage of *M. tuberculosis* infection, after the development of adaptive immunity. Consistent with reports in sterile inflammation models, we found that Ly6C^hi^ blood monocytes differentiate into multiple subsets of dendritic cells and macrophages in the lungs and lymph nodes of *M. tuberculosis*-infected mice. Using a recently-described method of distinguishing cells that reside in the vascular space from those in the tissue parenchyma, we determined that Ly6C^hi^ monocytes differentiate into Ly6C^lo^ monocytes that remain in the lung vascular space in *M. tuberculosis*-infected mice. Using the same method, we determined that differentiation of Ly6C^hi^ cells into other subsets happens after egress from the vascular compartment, without transitioning through a Ly6C^lo^ state. In particular, we determined that Ly6C^hi^ monocytes rapidly differentiate into Ly6C^hi^ CD11c^+^ MHCII^+^ population of DC, which is a transient intermediate state followed by differentiation into CD11b^+^ DC in the lungs. Although our evidence is indirect, we found kinetic evidence that both Ly6C^hi^ and CD11b^+^ DC migrate and transport bacteria from the lungs to the lung-draining mediastinal lymph node, consistent with previous reports [8-10].

We also determined that the response to *M. tuberculosis* infection in mice includes peripheral blood monocytosis, as observed in children and adult humans with TB [23-25], and using pulse-chase DNA labeling, we determined that peripheral blood monocytosis is due to enhanced production and egress from the bone marrow and not due to delayed egress from the blood. Pulse-chase DNA labeling further confirmed the capacity and kinetics of differentiation of monocyte-derived cells and established that the monocyte-derived professional phagocyte populations in the lungs and lymph nodes of *M. tuberculosis*-infected mice are dynamic and turn over with a half-life of less than one week. The finding that the monocyte-derived cell subsets that harbor *M. tuberculosis* turn over frequently suggested that the bacteria must exchange their cellular niches frequently, and we confirmed this prediction by detecting *M. tuberculosis* in newly-arrived cells in the lungs, indicating that the bacteria must respond regularly and rapidly to distinct cellular environments.

Although the dynamics of mononuclear cells in localized mycobacterial infections has gotten little recent attention, there are precedents in other animal models, and evidence to suggest similar events in humans. Nearly 50 years ago, administration of radiolabeled thymidine was used to characterize mononuclear cell recruitment and death at skin sites of injection of *M. bovis* BCG in rabbits, and revealed that in contrast to prior assumptions, granuloma mononuclear cell populations are dynamic [18, 36]. However, at the time those experiments were performed, it was not possible to distinguish distinct subsets of mononuclear cells or their differentiation other than by their morphology. More recent studies used adoptive transfer of bone marrow monocytes during *M. tuberculosis* infection of mice, and revealed tissue-dependent differentiation into multiple subsets of macrophages and dendritic cells, with distinct patterns of differentiation in the lungs and the lung-draining lymph nodes [15]. Our findings extend those results by revealing that Ly6C^hi^ CD11c^+^ cells appear after Ly6C^hi^ monocytes enter the lung parenchyma and are transient intermediates that subsequently differentiate into CD11c^hi^ CD11b^+^ dendritic cells. This observation helps reconcile apparently different results of analysis of the cells that transport *M. tuberculosis* from the lungs to the lung-draining mediastinal lymph node: one study attributed this property to inflammatory monocytes [8], while another reported that CD11b^+^ dendritic cells are responsible for bacterial transport [9]. Our present results indicate that the inflammatory monocytes credited with bacterial transport likely differentiate into Ly6C^hi^ CD11c^+^ dendritic cells after entering the lung parenchyma, where they acquire bacteria and differentiate further into CD11b^+^ dendritic cells, which contain bacteria and appear in the draining lymph node [9, 10].

Other recent studies have emphasized the importance of tissue-dependent differentiation of monocytes in tuberculosis, and certain of these have revealed important functional differences of monocyte-derived cell subsets. In particular, administration of poly-ICLC, a potent inducer of type I interferon expression, impaired control of *M. tuberculosis* in the lungs of mice, due to CCR2-dependent recruitment of monocyte-derived cells that are especially permissive for intracellular bacterial survival and/or growth [14]. In contrast, a recent study reported that interstitial macrophages that depended on CCL2 for their recruitment, are more able to constrain *M. tuberculosis* fitness as reflected by use of fluorescent reporter plasmids [2]. These seemingly contradictory results emphasize the potential plasticity of cells that differentiate from monocytes, which is likely the product of early programming of monocytes as well as of signals for differentiation and activation at the site of infection. Notably, studies in zebrafish embryos infected with virulent *M. marinum* have revealed that recruited macrophages are less able to control intracellular mycobacterial growth compared with tissue-resident macrophages [37]. As bacterial spread to recruited macrophages was a function of bacterial phenolic glycolipid, these results indicate that pathogen virulent factors can also act by directing bacteria to cells that are poorly equipped to kill them.

Although studies similar to ours have not been performed in humans or with human samples, it is noteworthy that cell subsets comparable to the ones studied here can be detected in human lung tissue [38]. Likewise, recent studies of human monocytes have revealed a comparable pattern of cell turnover and differentiation [39], indicating that in humans with tuberculosis, cell populations localized to the site of infection in granulomas may exhibit dynamic states of turnover and differentiation similar to those that we have found in *M. tuberculosis*-infected mice.

The finding that mononuclear cell populations are in a dynamic state during the chronic stage of *M. tuberculosis* infection may have clinical value. First, our findings that MNP populations are dynamic, even during the chronic phase of *M. tuberculosis* infection suggests that mechanisms that perturb the dynamics of MNP trafficking and differentiation may underlie or contribute to the transition from latent TB infection to active TB disease and/or the severity of TB disease. Evidence consistent with this hypothesis is that pulmonary TB in patients with diabetes mellitus, which has poorer outcomes compared with those of pulmonary TB in those without diabetes [40-42] is characterized by decreased frequencies of blood monocyte and dendritic cell populations compared with those in patients without diabetes [43]. Second, a recent study revealed that ‘trained hematopoietic stem cells’ that produce epigenetically-modified monocytes and macrophages are capable of contributing to control of *M. tuberculosis* infection in mice, in a manner that is independent of T lymphocytes [19]. Our finding that recruited macrophage populations are dynamic and that newly-arrived cells have access to bacteria even during the chronic stage of infection, implies that modification of cell populations to better restrict and/or kill *M. tuberculosis* may have therapeutic value in humans with drug-resistant tuberculosis, and may shorten the length of effective treatment, even in those with drug-susceptible tuberculosis.

## Methods

### Mice

C57/BL6 mice were bred in the New York University School of Medicine (New York, NY) animal facilities or purchased from The Jackson Laboratory and maintained under specific pathogen-free conditions. *Ptpcr^a^* (CD45.1) mice utilized in adoptive transfer experiments were either bred in the New York University School of Medicine animal facilities or purchased from Taconic Farms, Inc. Mice were infected with *M. tuberculosis* at 8-12 weeks of age. Uninfected or *M. tuberculosis*-infected mice were twice injected intraperitoneally with 2 mg of EdU (Thermo Fisher Scientific) in 400 ul of PBS, two hours apart.

### Ethics

All experiments were conducted in accordance with procedures approved by the NYU School of Medicine Institutional Animal Care and Use Committee and in accordance to the recommendations in the Guide for the Care and Use of Laboratory animals of the National Institutes of Health, operating under Animal Welfare Assurance number D16-00274. The protocol approval number was s15-01412.

### *M. tuberculosis* strains, growth and infection

*M. tuberculosis* (H37Rv) derivatives used to infect mice constitutively expressed either EGFP or mCherry. Bacteria were grown in Middlebrook 7H9 medium with 10% (v/v) albumin dextrose catalase (ADC) enrichment, 0.05% Tween80 and 50 µg/ml kanamycin or 25µg/ml hygromycin. Mice were infected via aerosol utilizing an inhalation exposure unit from Glas-Col. Mid-log cultures of *M. tuberculosis* were pelleted at 4000 g, resuspended in 7ml of PBS+0.05% Tween 80 and then serially centrifuged at 800 g to remove clumps. Clump-free cultures were then diluted to 1.5×10^6^ in dH20 and 5 mL of the inoculum was added to the nebulizer. Mice were infected with a program of 900s of preheating, 2400s of nebulization, 2400s of cloud decay, and 900s of decontamination. For controls of fluorescent-protein-expressing *M. tuberculosis*, mice were infected with wild-type H37Rv by the same procedure on the same day. Infection dose was determined by euthanizing mice within 24 h of infection, plating lung homogenates on 7H11 agar plates, and counting CFU within 2-3 weeks of incubation at 37° C.

### Tissue harvests and processing

Mice were anesthetized by inhalation of 10% Isoflurane in mineral oil (v/v) and then retro-orbitally injected with 2.5 µg of anti-CD45-APC-Cy7 antibody (clone 30-F11, Biolegend) in 200 µl of PBS. Three minutes after antibody injection mice were euthanized by Isoflurane inhalation and cervical dislocation. Lungs, MLN, peripheral LN, BM and blood were collected from the mice depending on the experiment. Solid tissues were placed in complete media (RPMI1640, 5% FCS, 10 mM HEPES, 1x NEAA, 1 mM Sodium Pyruvate, 55 µM 2-mercaptoethanol) containing 1 mg/ml type 2 collagenase and 30 µg/ml DNase, minced with scissors and incubated for 30 minutes at 37° C. Tissues were then forced through a 70-uM cell strainer (BD) and washed twice in collagenase wash buffer (PBS + 2% FBS (v/v), 2 mM EDTA). Mouse femur and tibia leg bones were cleaned of tissue before removing their ends with scissors. Marrow was then extracted by perfusing with 5 mL of complete media using a 27G needle. Blood was collected in complete media. RBC were removed from all tissues using ACK lysis buffer (155 mM NH_3_Cl, 10 mM KHCO_3_, and 88 µM EDTA) and live cells were counted using trypan blue exclusion and a Countess cell counter (Thermo Fischer Scientific)

### Flow cytometry

Single cell suspensions were first washed with PBS and stained Zombie-Aqua (Biolegend) live/dead dye for 20 min at 4°C. Cells were then incubated with FcγRIII/I blocking antibody (clone 2.4G2)(Biolegend) and fluorescently labeled antibodies in PBS + 3% BSA (w/v) for 30 minutes at 4° C. From Biolegend, anti-CD11b (clone M1/70), CD11c (N418), MHCII I-A and I-E (M5/114.15.2), CD19 (6D5), Ly6C (AL-21), Ly6G (1A8), NK1.1 (PK136), Thy1.2 (30-H12), and CD103 (2E7) or XCR1 (ZET). From BD, anti-B220 (RA3-6B2), CD45 (30-F11) and Siglec F (E50-2440). Cells were then washed twice with PBS+3% FCS and fixed overnight in 1% paraformaldehyde (v/v).

For EdU staining, cell suspensions were stained with live/dead dye and incubated with blocking and fluorescently-labeled antibodies, except for antibodies with PE or PE-conjugated fluorophores. Cells were washed and fixed in 4% paraformaldehyde before permeablization in Click-iT saponin-based perm-wash buffer (Molecular Probes). EdU was then labeled with Alexa Fluor 647 azide in Click-iT buffer as per the manufacturers protocol. After further washing, cells were then incubated with antibodies labeled with PE and PE-conjugates. Cells were then washed twice with PBS+3% FCS and fixed overnight in 1% paraformaldehyde (v/v). Samples were acquired on a BD Biosciences LSR II and analyzed using FlowJo (TreeStar, Inc.).

### Immunofluorescence

Mice infected with GFP-expressing *M. tuberculosis* were injected with EdU and euthanized as described. Lungs were instilled with OCT by syringe via the trachea prior to embedding in OCT and freezing in liquid N_2_. Frozen tissues were then cut into 6uM with a cryostat, mounted on slides and fixed immediately with ice-cold 100% acetone for 15 minutes then stored at −20° C. Slides were rehydrated in PBS then blocked for PBS + 5% FBS for 15 min. EdU was labeled with Alexa Fluor 647 in Click-iT buffer as per the manufacturers protocol. Tissue sections were blocked with anti-FcγRIII/I (2.4G2) and labeled with anti-CD11b and CD4 fluorescently conjugated antibodies. Slides were mounted with Prolong-Diamond (Life sciences). Images were captured using the 60x oil-immersion objective on a Nikon Eclipse Ti fluorescent microscope. Images were captured over multiple frames and z-stacks and were stitched together and deconvoluted using NIS Elements software (Nikon). Images were further analyzed using ImageJ (National Institutes of Health).

### Adoptive transfer of monocytes

Bone marrow and blood were collected from 30 naïve CD45.1 mice and processed into single cell suspensions. Cells were incubated with biotin-conjugated Monocyte Isolation Kit antibodies (Miltenyi Biotec) and anti-biotin microbeads as per the manufacturers instructions. Monocytes were then enriched negative selection on either an AutoMACS or MultiMACS magnetic column. Stem cells were further depleted by incubating enriched monocytes with biotin-conjugated Sca-1 (D7), CD117 (2B8) and CD135 (A2F10) antibodies and anti-biotin microbeads. Monocytes were then washed twice in PBS, counted and 1.0-2.25 × 10^6^ monocytes in 100 µl of PBS were retro-orbitally intravenously injected into each *M. tuberculosis*-infected mouse.

### CD62L blockade

Mice infected with aerosolized *M. tuberculosis* were intravenously injected with 250 µg of isotype (clone 2A3) or anti-CD62L antibody (Mel-14)(BioXCell) on day 28 post infection. Two hours after administering antibodies, mice were intraperitoneally injected with 1 mg of EdU. Mice were given an additional 100 µg of isotype or anti-CD62L antibody by intraperitoneal injection 24 h after the first antibody injection. Two days after the first antibody treatment, mice were injected with anti-CD45 antibody, sacrificed and organs were collected and processed as described previously.

## Acknowledgments

We thank Amber Cornelius, Miriam Bolz, and Jessica Jang for their contributions to the success of this work, and to members of the Ernst, Philips, and Portal laboratories for spirited discussions.

## Supporting Information

**S1 Fig. Gating hierarchy for multiple professional phagocyte subsets in the lungs of *M. tuberculosis*-infected mice.** Representative gating strategy to identify mononuclear cell subsets within the lungs of *M. tuberculosis*-infected mice.

**S2 Fig. Kinetics of Ly6C^hi^ monocyte turnover during *M. tuberculosis* infection.** Bone marrow monocytes (CD45.1^+^) were depleted of stem cells and transferred to *M. tuberculosis*-infected mice (CD45.2^+^). (A) Representative plots of CD45.1 donor-derived mononuclear cell subsets in the lungs of mice 40 hours after transfer to *M. tuberculosis*-infected mice. (B) The proportion of Ly6C^lo^CD11b^+^CD11c^lo^MHCII^−^ cells that are within the parenchyma increases as *M. tuberculosis* infection increases. Data are from 1-4 experiments per infection phase with 29-30 mice per week per experiment. (C) Mean fluorescent intensity (MFI) of professional phagocyte populations in the lungs of mice infected for 8 weeks with *M. tuberculosis*. Data are calculated by subtracting the background MFI of fluorescence minus one (FMO) control samples from the PU.1 MFI and are presented as the mean ± SEM of pooled samples from 5 mice and 4 time points of a single experiment. (D) Calculated half-life of donor total donor CD45.1 cells or Ly6C^hi^ monocytes and R^2^ coefficient of determination. (E) Comparison of fits for the slopes of the best-fit lines for total donor cells and donor Ly6C^hi^ monocytes in each tissue. The null hypothesis is that all slopes are equal, and each equation is compared to a best-fit equation calculated from all data points. Data from D and E are from a single representative experiment of two experiments and are presented as means ± SEM of 5 mice per time point.

**S3 Fig. Turnover of recently-proliferated Ly6C^hi^ monocytes during *M*. *tuberculosis* infection.** Monocytosis in the blood of *M. tuberculosis*-infected mice is matched by an increase in the rate of Ly6C^hi^ monocyte turnover. (A) Peripheral blood monocytosis during chronic *M. tuberculosis* infection. The Fig shows the total numbers of Ly6C^hi^ monocytes in the blood of uninfected and *M. tuberculosis*-infected mice, by time post-infection. Data are presented as blood monocytes in individual mice from 1-4 experiments per infection phase with 29-30 mice per week, per experiment. (B) Frequency of EdU^+^ staining in Ly6C^hi^ monocytes in multiple tissues of uninfected and *M. tuberculosis-*infected mice and the exponential best-fit line. (C) Total numbers of EdU^+^ Ly6C^hi^ monocytes in multiple tissues of uninfected and *M. tuberculosis*infected mice and the exponential best-fit line. Data are means from 1-4 experiments per infection phase with 4-5 mice per time point per experiment.

**S1 Table Half-lives of Ly6C^hi^ monocytes during multiple phases of *M. tuberculosis* infection.** *M. tuberculosis*-infected mice were injected with EdU and its incorporation by dividing monocytes was evaluated by flow cytometry at multiple time points. Exponential one-phase decay calculations were performed for Ly6C^hi^ monocytes during each infection phase and in each tissue. (A) Tabulated results of half-lives calculated from exponential best-fit lines of %EdU^+^ Ly6C^hi^ monocytes and the R^2^ coefficient of determination. Comparison of fits for the slopes of the best-fit lines of %EdU^+^ Ly6C^hi^ monocytes for each organ from infected or naïve mice. Null hypothesis is that the slope is the same for all each tissue, independent of infection phase. (B) Tabulated results of half-lives calculated from exponential best-fit lines of the total number of EdU^+^ Ly6C^hi^ monocytes and the R^2^ coefficient of determination. Comparison of fits for the slopes of the best-fit lines of total number of EdU^+^ Ly6C^hi^ monocytes for each organ from infected or naïve mice. Null hypothesis is that the slope is the same for all each tissue, independent of infection phase. Calculated best-fit of all infection phases for each organ and the R^2^ coefficient of determination.

**S2 Table Mean fold increase in lung mononuclear cell populations relative to uninfected mice.** Mean fold-increase of lung mononuclear cell populations at multiple phases of *M. tuberculosis* infection, relative to uninfected mice. Data are means from 1-4 experiments per infection phase with 4-5 mice per time point per experiment.

**S4 Fig. Composition of mononuclear cell subsets at multiple phases of *M. tuberculosis* infection.** *M. tuberculosis*-infected mice were injected with EdU and its incorporation by mononuclear cell subsets was evaluated by flow cytometry at multiple time points. (A) Total numbers, frequency EdU^+^ (B) and total numbers (C) of EdU^+^ peripheral blood and lung vasculature Ly6C^lo^ monocytes in uninfected mice or mice pulsed with EdU 4 weeks, 8 weeks and 16 weeks after infection with *M. tuberculosis.* Data are means and SEM from 1-4 experiments per infection phase with 4-5 mice per time point per experiment. (D) Mean percent of maximum frequency-of-live cells for EdU^+^ Ly6C^lo^ monocytes in the lung vasculature and circulation of uninfected mice or mice pulsed with EdU 4 weeks, 8 weeks and 16 weeks after infection with *M. tuberculosis.* (E) Total numbers and relative frequency (F) of CD45 IV^−^ professional phagocytes in the lung parenchyma of *M. tuberculosis*-infected mice at multiple phases of infection. (G) Frequency of Ki67^+^ staining on multiple professional phagocytes from individual mice in the lung parenchyma 4 weeks after *M. tuberculosis* infection. Data are presented as individual mice from a single experiment. (H) Total area under the curve of vascular and parenchymal EdU^+^ mononuclear cell subsets in the lungs of uninfected mice or *M. tuberculosis*-infected mice pulsed with EdU at weeks 4, 8 and 16 post infection. Data are means and SEM from 1-4 experiments per infection phase with 4-5 mice per time point per experiment.

**S5 Fig. MLN mononuclear cell subsets proliferate locally or are derived from lung cells, not the blood.** *M. tuberculosis*-infected mice were injected with EdU and its incorporation by mononuclear cell subsets in the MLN was evaluated by flow cytometry at multiple time points. (A) Representative gating hierarchy for multiple mononuclear cell subsets within the MLN of mice infected with *M. tuberculosis*. (B) Relative frequency of professional phagocytes containing mCherry-expressing *M. tuberculosis* in the MLN of *M. tuberculosis*-infected mice at multiple phases of infection. Data are presented as means ± SEM of 1-3 experiments with 5 mice per time point. (C) Schematic showing the experimental design to block monocyte trafficking to the LN through the HEV. (D) Total numbers of Ly6C^hi^ monocytes in multiple tissues and in the blood in mice day 30 post-infection and following two days of isotype or anti-CD62L antibody treatment. (E) Frequency of EdU incorporation by Ly6C^hi^ monocytes in multiple tissues and the blood two days following EdU pulse of week 4 *M. tuberculosis*-infected mice. Data from D and E are presented as means ± SEM from 2 experiments with 4-5 mice per time point per experiment. Statistics are 2-way ANOVA with Sidak’s multiple comparisons tests. *p<0.05

**S6 Fig. *M. tuberculosis* infection of recently-proliferated neutrophils and mononuclear cells**. Mice infected with fluorescent protein-expressing *M. tuberculosis* were injected with EdU and its incorporation by dividing myeloid cells evaluated by flow cytometry at multiple time points. (A) Frequency of EdU^+^ neutrophils in the lung vasculature and parenchyma of uninfected mice or mice pulsed with EdU 4 weeks, 8 weeks and 16 weeks after infection with *M. tuberculosis*. Data are presented as means and SEM from 1-4 experiments per infection phase with 4-5 mice per time point per experiment. (B) Composition of total lung cells infected with mCherry-expressing *M. tuberculosis* at multiple phases of infection. Data are presented as means and SEM from 1-4 experiments with 5 mice per time point. (C) Frequency of Rv^+^ cells within EdU^+^ mononuclear cells in the lung parenchyma of mice pulsed with EdU 16 weeks after infection with *M. tuberculosis*. (D) Total numbers of Rv^+^ EdU^+^ mononuclear cells in the lung parenchyma of mice pulsed with EdU 16 weeks after infection with *M. tuberculosis*. (E) Flow cytometric analysis of the frequency of Rv-mCherry-infected professional phagocytes that were EdU^+^ at multiple time points following EdU pulse 16 weeks post infection. Data are presented as means ± SEM of 4-5 mice per time point from a single infection for each phase. Statistics are 2-way ANOVA with Sidak’s multiple comparisons tests. *p<0.05, **p<0.01, ***p<0.001, ****p<0.0001.

## References

1. Mwandumba HC, Russell DG, Nyirenda MH, Anderson J, White SA, Molyneux ME, et al. Mycobacterium tuberculosis resides in nonacidified vacuoles in endocytically competent alveolar macrophages from patients with tuberculosis and HIV infection. Journal of immunology. 2004;172(7):4592–8. Epub 2004/03/23. PubMed PMID: 15034077.

2. Huang L, Nazarova EV, Tan S, Liu Y, Russell DG. Growth of Mycobacterium tuberculosis in vivo segregates with host macrophage metabolism and ontogeny. The Journal of experimental medicine. 2018. doi: 10.1084/jem.20172020. PubMed PMID: 29500179.

3. Sabin FR, Doan CA. The Relation of Monocytes and Clasmatocytes to Early Infection in Rabbits with Bovine Tubercle Bacilli. The Journal of experimental medicine. 1927;46(4):627–44. PubMed PMID: 19869362; PubMed Central PMCID: PMC2131307.

4. Wolf AJ, Linas B, Trevejo-Nunez GJ, Kincaid E, Tamura T, Takatsu K, et al. Mycobacterium tuberculosis infects dendritic cells with high frequency and impairs their function in vivo. Journal of immunology. 2007;179(4):2509–19. PubMed PMID: 17675513.

5. Davis JM, Ramakrishnan L. The role of the granuloma in expansion and dissemination of early tuberculous infection. Cell. 2009;136(1):37–49. doi: 10.1016/j.cell.2008.11.014. PubMed PMID: 19135887; PubMed Central PMCID: PMC3134310.

6. Pagan AJ, Ramakrishnan L. The Formation and Function of Granulomas. Annual review of immunology. 2018. Epub 2018/02/06. doi: 10.1146/annurev-immunol-032712-100022. PubMed PMID: 29400999.

7. Tailleux L, Schwartz O, Herrmann JL, Pivert E, Jackson M, Amara A, et al. DC-SIGN is the major Mycobacterium tuberculosis receptor on human dendritic cells. The Journal of experimental medicine. 2003;197(1):121–7. Epub 2003/01/08. PubMed PMID: 12515819; PubMed Central PMCID: PMCPMC2193794.

8. Samstein M, Schreiber HA, Leiner IM, Susac B, Glickman MS, Pamer EG. Essential yet limited role for CCR2(+) inflammatory monocytes during Mycobacterium tuberculosis-specific T cell priming. eLife. 2013;2:e01086. doi: 10.7554/eLife.01086. PubMed PMID: 24220507; PubMed Central PMCID: PMC3820971.

9. Srivastava S, Ernst JD. Cell-to-cell transfer of M. tuberculosis antigens optimizes CD4 T cell priming. Cell host & microbe. 2014;15(6):741–52. doi: 10.1016/j.chom.2014.05.007. PubMed PMID: 24922576; PubMed Central PMCID: PMC4098643.

10. Wolf AJ, Desvignes L, Linas B, Banaiee N, Tamura T, Takatsu K, et al. Initiation of the adaptive immune response to Mycobacterium tuberculosis depends on antigen production in the local lymph node, not the lungs. The Journal of experimental medicine. 2008;205(1):105–15. doi: 10.1084/jem.20071367. PubMed PMID: 18158321; PubMed Central PMCID: PMC2234384.

11. Srivastava S, Grace PS, Ernst JD. Antigen Export Reduces Antigen Presentation and Limits T Cell Control of M. tuberculosis. Cell host & microbe. 2016;19(1):44–54. doi: 10.1016/j.chom.2015.12.003. PubMed PMID: 26764596; PubMed Central PMCID: PMC4715867.

12. Yona S, Kim KW, Wolf Y, Mildner A, Varol D, Breker M, et al. Fate mapping reveals origins and dynamics of monocytes and tissue macrophages under homeostasis. Immunity. 2013;38(1):79–91. doi: 10.1016/j.immuni.2012.12.001. PubMed PMID: 23273845; PubMed Central PMCID: PMC3908543.

13. Landsman L, Varol C, Jung S. Distinct differentiation potential of blood monocyte subsets in the lung. Journal of immunology. 2007;178(4):2000–7. Epub 2007/02/06. PubMed PMID: 17277103.

14. Antonelli LR, Gigliotti Rothfuchs A, Goncalves R, Roffe E, Cheever AW, Bafica A, et al. Intranasal Poly-IC treatment exacerbates tuberculosis in mice through the pulmonary recruitment of a pathogen-permissive monocyte/macrophage population. The Journal of clinical investigation. 2010;120(5):1674–82. Epub 2010/04/15. doi: 10.1172/jci40817. PubMed PMID: 20389020; PubMed Central PMCID: PMCPMC2860920.

15. Skold M, Behar SM. Tuberculosis triggers a tissue-dependent program of differentiation and acquisition of effector functions by circulating monocytes. Journal of immunology. 2008;181(9):6349–60. PubMed PMID: 18941226.

16. Martin CJ, Booty MG, Rosebrock TR, Nunes-Alves C, Desjardins DM, Keren I, et al. Efferocytosis is an innate antibacterial mechanism. Cell host & microbe. 2012;12(3):289–300. Epub 2012/09/18. doi: 10.1016/j.chom.2012.06.010. PubMed PMID: 22980326; PubMed Central PMCID: PMCPMC3517204.

17. Repasy T, Martinez N, Lee J, West K, Li W, Kornfeld H. Bacillary replication and macrophage necrosis are determinants of neutrophil recruitment in tuberculosis. Microbes and infection / Institut Pasteur. 2015;17(8):564–74. Epub 2015/04/12. doi: 10.1016/j.micinf.2015.03.013. PubMed PMID: 25862076; PubMed Central PMCID: PMCPMC4508221.

18. Dannenberg AM, Jr., Ando M, Shima K. Macrophage accumulation, division, maturation, and digestive and microbicidal capacities in tuberculous lesions. 3. The turnover of macrophages and its relation to their activation and antimicrobial immunity in primary BCG lesions and those of reinfection. Journal of immunology. 1972;109(5):1109–21. PubMed PMID: 5079091.

19. Kaufmann E, Sanz J, Dunn JL, Khan N, Mendonca LE, Pacis A, et al. BCG Educates Hematopoietic Stem Cells to Generate Protective Innate Immunity against Tuberculosis. Cell. 2018;172(1-2):176–90.e19. Epub 2018/01/13. doi: 10.1016/j.cell.2017.12.031. PubMed PMID: 29328912.

20. Carlin LM, Stamatiades EG, Auffray C, Hanna RN, Glover L, Vizcay-Barrena G, et al. Nr4a1-dependent Ly6C(low) monocytes monitor endothelial cells and orchestrate their disposal. Cell. 2013;153(2):362–75. Epub 2013/04/16. doi: 10.1016/j.cell.2013.03.010. PubMed PMID: 23582326; PubMed Central PMCID: PMCPMC3898614.

21. Hanna RN, Carlin LM, Hubbeling HG, Nackiewicz D, Green AM, Punt JA, et al. The transcription factor NR4A1 (Nur77) controls bone marrow differentiation and the survival of Ly6C- monocytes. Nat Immunol. 2011;12(8):778–85. Epub 2011/07/05. doi: 10.1038/ni.2063. PubMed PMID: 21725321; PubMed Central PMCID: PMCPMC3324395.

22. Menezes S, Melandri D, Anselmi G, Perchet T, Loschko J, Dubrot J, et al. The Heterogeneity of Ly6C(hi) Monocytes Controls Their Differentiation into iNOS(+) Macrophages or Monocyte-Derived Dendritic Cells. Immunity. 2016;45(6):1205–18. Epub 2016/12/22. doi: 10.1016/j.immuni.2016.12.001. PubMed PMID: 28002729; PubMed Central PMCID: PMCPMC5196026.

23. Rogers PM. A Study of the Blood Monocytes in Children with Tuberculosis. New England Journal of Medicine. 1928;198(14):740–9. doi: 10.1056/nejm192805241981410.

24. La Manna MP, Orlando V, Dieli F, Di Carlo P, Cascio A, Cuzzi G, et al. Quantitative and qualitative profiles of circulating monocytes may help identifying tuberculosis infection and disease stages. PloS one. 2017;12(2):e0171358. doi: 10.1371/journal.pone.0171358.

25. Naranbhai V, Hill AVS, Abdool Karim SS, Naidoo K, Abdool Karim Q, Warimwe GM, et al. Ratio of Monocytes to Lymphocytes in Peripheral Blood Identifies Adults at Risk of Incident Tuberculosis Among HIV-Infected Adults Initiating Antiretroviral Therapy. The Journal of infectious diseases. 2014;209(4):500–9. doi: 10.1093/infdis/jit494.

26. Ginhoux F, Liu K, Helft J, Bogunovic M, Greter M, Hashimoto D, et al. The origin and development of nonlymphoid tissue CD103+ DCs. The Journal of experimental medicine. 2009;206(13):3115–30. doi: 10.1084/jem.20091756. PubMed PMID: 20008528; PubMed Central PMCID: PMC2806447.

27. Becker M, Guttler S, Bachem A, Hartung E, Mora A, Jakel A, et al. Ontogenic, Phenotypic, and Functional Characterization of XCR1(+) Dendritic Cells Leads to a Consistent Classification of Intestinal Dendritic Cells Based on the Expression of XCR1 and SIRPalpha. Front Immunol. 2014;5:326. Epub 2014/08/15. doi: 10.3389/fimmu.2014.00326. PubMed PMID: 25120540; PubMed Central PMCID: PMCPMC4112810.

28. Bachem A, Hartung E, Guttler S, Mora A, Zhou X, Hegemann A, et al. Expression of XCR1 Characterizes the Batf3-Dependent Lineage of Dendritic Cells Capable of Antigen Cross-Presentation. Front Immunol. 2012;3:214. Epub 2012/07/25. doi: 10.3389/fimmu.2012.00214. PubMed PMID: 22826713; PubMed Central PMCID: PMCPMC3399095.

29. Jakubzick C, Gautier EL, Gibbings SL, Sojka DK, Schlitzer A, Johnson TE, et al. Minimal differentiation of classical monocytes as they survey steady-state tissues and transport antigen to lymph nodes. Immunity. 2013;39(3):599–610. doi: 10.1016/j.immuni.2013.08.007. PubMed PMID: 24012416; PubMed Central PMCID: PMC3820017.

30. Blomgran R, Ernst JD. Lung neutrophils facilitate activation of naive antigen-specific CD4+ T cells during Mycobacterium tuberculosis infection. Journal of immunology. 2011;186(12):7110–9. doi: 10.4049/jimmunol.1100001. PubMed PMID: 21555529; PubMed Central PMCID: PMC3376160.

31. Chackerian AA, Alt JM, Perera TV, Dascher CC, Behar SM. Dissemination of Mycobacterium tuberculosis is influenced by host factors and precedes the initiation of T-cell immunity. Infection and immunity. 2002;70(8):4501–9. PubMed PMID: 12117962; PubMed Central PMCID: PMCPMC128141.

32. Behr MA, Waters WR. Is tuberculosis a lymphatic disease with a pulmonary portal? The Lancet Infectious diseases. 2014;14(3):250–5. Epub 2013/11/26. doi: 10.1016/s1473-3099(13)70253-6. PubMed PMID: 24268591.

33. Harakawa N, Shigeta A, Wato M, Merrill-Skoloff G, Furie BC, Furie B, et al. P-selectin glycoprotein ligand-1 mediates L-selectin-independent leukocyte rolling in high endothelial venules of peripheral lymph nodes. International immunology. 2007;19(3):321–9. doi: 10.1093/intimm/dxl149. PubMed PMID: 17267415.

34. Leon B, Ardavin C. Monocyte migration to inflamed skin and lymph nodes is differentially controlled by L-selectin and PSGL-1. Blood. 2008;111(6):3126–30. doi: 10.1182/blood-2007-07-100610. PubMed PMID: 18184867.

35. Xu H, Manivannan A, Crane I, Dawson R, Liversidge J. Critical but divergent roles for CD62L and CD44 in directing blood monocyte trafficking in vivo during inflammation. Blood. 2008;112(4):1166–74. doi: 10.1182/blood-2007-06-098327. PubMed PMID: 18391078; PubMed Central PMCID: PMC2515150.

36. Ando M, Dannenberg AM, Jr. Macrophage accumulation, division, maturation, and digestive and microbicidal capacities in tuberculous lesions. IV. Macrophage turnover, lysosomal enzymes, and division in healing lesions. Laboratory investigation; a journal of technical methods and pathology. 1972;27(5):466–72. PubMed PMID: 4569599.

37. Cambier CJ, O’Leary SM, O’Sullivan MP, Keane J, Ramakrishnan L. Phenolic Glycolipid Facilitates Mycobacterial Escape from Microbicidal Tissue-Resident Macrophages. Immunity. 2017;47(3):552–65.e4. Epub 2017/08/29. doi: 10.1016/j.immuni.2017.08.003. PubMed PMID: 28844797; PubMed Central PMCID: PMCPMC5610147.

38. Yu YR, Hotten DF, Malakhau Y, Volker E, Ghio AJ, Noble PW, et al. Flow Cytometric Analysis of Myeloid Cells in Human Blood, Bronchoalveolar Lavage, and Lung Tissues. American journal of respiratory cell and molecular biology. 2016;54(1):13–24. Epub 2015/08/13. doi: 10.1165/rcmb.2015-0146OC. PubMed PMID: 26267148; PubMed Central PMCID: PMCPMC4742930.

39. Patel AA, Zhang Y, Fullerton JN, Boelen L, Rongvaux A, Maini AA, et al. The fate and lifespan of human monocyte subsets in steady state and systemic inflammation. The Journal of experimental medicine. 2017;214(7):1913–23. Epub 2017/06/14. doi: 10.1084/jem.20170355. PubMed PMID: 28606987; PubMed Central PMCID: PMCPMC5502436.

40. Faurholt-Jepsen D, Range N, PrayGod G, Jeremiah K, Faurholt-Jepsen M, Aabye MG, et al. Diabetes is a strong predictor of mortality during tuberculosis treatment: a prospective cohort study among tuberculosis patients from Mwanza, Tanzania. Tropical medicine & international health: TM & IH. 2013;18(7):822–9. Epub 2013/05/08. doi: 10.1111/tmi.12120. PubMed PMID: 23648145.

41. Gil-Santana L, Almeida-Junior JL, Oliveira CA, Hickson LS, Daltro C, Castro S, et al. Diabetes Is Associated with Worse Clinical Presentation in Tuberculosis Patients from Brazil: A Retrospective Cohort Study. PloS one. 2016;11(1):e0146876. Epub 2016/01/12. doi: 10.1371/journal.pone.0146876. PubMed PMID: 26752596; PubMed Central PMCID: PMCPMC4709051.

42. Jimenez-Corona ME, Cruz-Hervert LP, Garcia-Garcia L, Ferreyra-Reyes L, Delgado-Sanchez G, Bobadilla-Del-Valle M, et al. Association of diabetes and tuberculosis: impact on treatment and post-treatment outcomes. Thorax. 2013;68(3):214–20. Epub 2012/12/20. doi: 10.1136/thoraxjnl-2012-201756. PubMed PMID: 23250998; PubMed Central PMCID: PMCPMC3585483.

43. Kumar NP, Moideen K, Sivakumar S, Menon PA, Viswanathan V, Kornfeld H, et al. Modulation of dendritic cell and monocyte subsets in tuberculosis-diabetes co-morbidity upon standard tuberculosis treatment. Tuberculosis. 2016;101:191–200. Epub 2016/11/21. doi: 10.1016/j.tube.2016.10.004. PubMed PMID: 27865391; PubMed Central PMCID: PMCPMC5127284.

